# Biochemical characterization of AniA from *Neisseria gonorrhoeae*

**DOI:** 10.1101/2022.04.07.487406

**Authors:** Daniela S. Barreiro, Ricardo N. S. Oliveira, Sofia R. Pauleta

## Abstract

AniA, the nitrite reductase from *Neisseria gonorrhoeae*, has been shown to play a crucial role in the infection mechanism of this microorganism by producing NO and abolishing epithelial exfoliation. This enzyme is a trimer with one type-1 copper center per subunit and one type 2 copper center in the subunits interface, with the latter being the catalytic site. The two centers were characterized by visible, EPR and CD spectroscopy for the first time, indicating that AniA’s type 1 copper center has a high rhombicity, which is attributed to its tetrahedral geometry, and shorter Met-Cu bond, while type 2 copper center has the usual properties, though with a shorter hyperfine coupling constant (A//= 9.1 mT). The thermostability of AniA was analyzed by differential scanning calorimetry showing a single endothermic transition in the thermogram, with a maximum at 95 °C, while the CD spectra in the visible region indicates the presence of copper centers at 85-90 °C. The reoxidation rates of AniA in the presence of nitrite were analyzed by visible spectroscopy showing a pH dependence and being higher at pH 6.0. The high thermostability of this enzyme might be important for maintaining a high activity in the extracellular space and be less prone to denaturation and proteolysis, contributing to the proliferation of *N. gonorrhoeae*.

## 1. Introduction

Nitrite reductases (NiR) are copper or heme-containing enzymes that catalyze the reduction of nitrite (NO_2_^-^) to nitric oxide (NO) in the periplasm of gram-negative bacteria [1], according to equation 1, as the second step of the denitrification pathway.

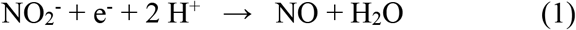

The heme-containing enzymes, *cd*_1_NiR, are homodimers, with each subunit containing one *d*_1_-type heme and one *c*-type heme, with the first being the catalytic site, and the second being responsible for receiving electrons from small redox proteins, such as *c*-type cytochromes or cupredoxins, and channeling them to the active site [2-4].

The copper-containing enzymes, *cd*_1_NiR, have two copper centers with similar functions. The electrons enter through the type 1 copper center (T1Cu) and are then transferred to the type 2 copper center (T2Cu), where NO_2_^-^ is converted to NO [5-9].

In recent years with the advances in genome sequencing, the gene encoding CuNiRs has been shown to be associated with additional domains either at the N- or at the C-terminus. These additional domains bind either *c*-type hemes or T1Cu centers and mediate electron transfer to the core unit [10]. This core unit forms a homotrimeric structure, with each subunit having two copper centers, one T1Cu and one T2Cu (Figure 1), that have different structural, spectroscopic, and functional properties [11]. T1Cu copper is coordinated by two histidine residues, a methionine, and a cysteine, from the same subunit, and connects to the T2Cu through a histidine-cysteine bridge. T2Cu, the active site, is coordinated by three histidine residues, two of which from the other subunit being position at the interface between subunits (Figure 1) [12]. Both centers are connected through a cysteine-histidine bridge. Other residues in the catalytic pocket were reported to play an important role in the catalytic cycle, such as the conserved aspartate residue (Asp137, Figure 1) [13-15] involved in the formation of T2Cu^+^-NO_2_^-^ intermediate species, by triggering the electron transfer through the cysteine-histidine bridge, thus acting as a “substrate-sensing pathway” [7, 8, 16, 17].

**Figure 1.**
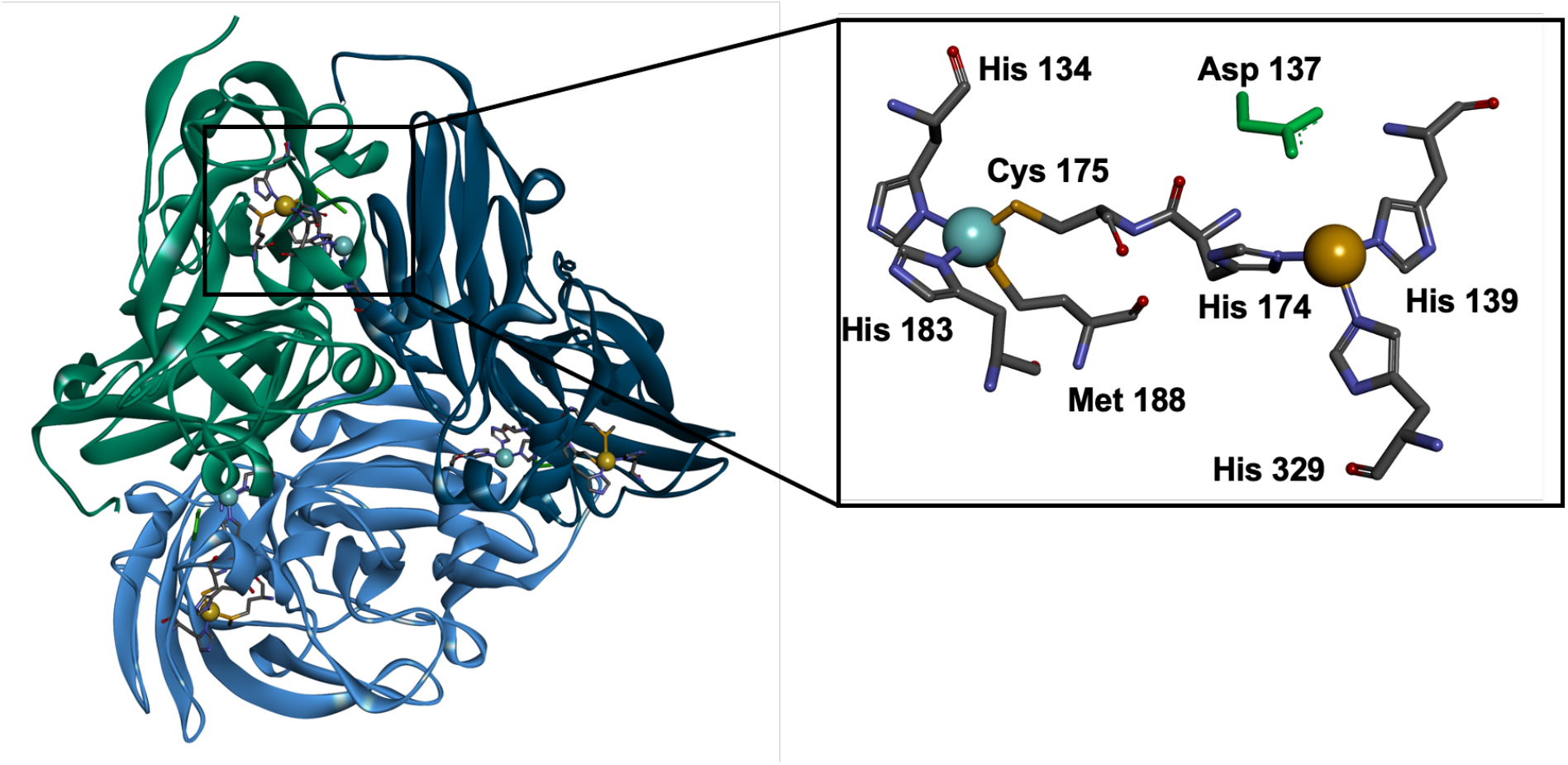
Structure of AniA from *Neisseria gonorrhoeae* as a trimer, evidencing the location of T1Cu center (blue sphere) in each subunit and T2Cu center (brown sphere) in the interface between the subunits. The coordination of the two centers is shown on the left panel. Figures were prepared in Discovery Studio Vizualizer using the coordinates PDB ID 1KBW.

CuNiRs can be divided into three classes, blue, green and bluish-green, depending on the spectroscopic properties of their T1Cu center [11], as T2Cu center has no absorption bands in the visible region. The typical absorption band of T1Cu center has a maximum at around 600 nm, corresponding to a S (π) Cys→Cu charge transfer transition [18]. The CuNiR with such spectral features belong to the blue CuNiR. However, most enzymes belong to the bluish-green classes and have an additional absorption band at around 460 nm attributed to a S (σ) Cys→Cu charge transfer transition. In the absorption spectra of green CuNiR, besides the low intensity absorption band at 600 nm and a more intense band centered at 450 nm, have a shoulder at 400 nm that has been assigned at a S(δ) Met→Cu charge transfer transition. The absorbance ratio between these two bands is a consequence of T1Cu geometry and length of the bond between the copper atom and sulfur of the methionine and cysteine side chain [18].

*Neisseria gonorrhoeae* is an obligate human pathogen responsible for the sexually transmitted disease gonorrhea, which affects millions of people worldwide. As of today, no effective vaccine against *N. gonorrhoeae* is available, and resistant strains against the used antibiotics is increasing, making the study of its survival mechanisms important as they constitute good drug targets. One of these mechanisms in the anaerobic respiration. *N. gonorrhoeae* possesses a truncated denitrification pathway, comprised by a nitrite reductase, AniA, and a nitric oxide reductase, qNorB, that allows the use of N-based compounds as alternative electron acceptors for energy conservation [19]. In addition, considering that NO is a modulating molecule of the immune response, as low levels (nM) of NO are anti-inflammatory and high levels of NO (μM) are pro-inflammatory [20], qNOR will be able to lower these levels. Moreover, nitrite reductase has been shown to play an important role in infection by producing NO and initiating a eukaryotic signaling pathway that leads to suppression of epithelial exfoliation [21, 22].

AniA is a copper nitrite reductase, CuNiR, belonging to the bluish-green subclass. It is described as an homotrimer and is found in the periplasm of *N gonorrhoeae*, anchored to the outer membrane by a palmitate bound to a cysteine residue [23]. The redox partner of this enzyme is still not known and was shown not to be the lipid-modified azurin from the same organism [24].

In here we report the first stability study using differential scanning calorimetry and circular dichroism, which indicated that this is a highly stable enzyme. In addition, its spectroscopic properties are described for the first time. Given the importance of this enzyme during infection it constitutes an excellent target to the design of novel therapies and thus the knowledge about its stability and spectroscopic can help identify compounds that either destabilize its structure or interact with the active site.

## 2. Materials and Methods

### 2.1 Heterologous Production of AniA

AniA was heterologously produced in *Escherichia coli* BL21(DE3) competent cells (Merck). The expression vector encoding the soluble domain of AniA (pET28A_*aniA*) was a kind gift from Prof. Michael E. P. Murphy (from the University of British Columbia, Canada) using the cloning protocol of *aniA* gene described in [15]. Four to five colonies of *E. coli* BL21(DE3) cells transformed with pET28a_AniA were selected to inoculate 50 mL of Luria-Bertani medium, supplemented with 30 μg/mL of kanamycin sulfate and grown overnight at 37 °C, 210 rpm. Fresh 2xYT medium, supplemented with 30 μg/mL of kanamycin and 0.1 mM of CuSO_4_, was inoculated with 2 % of the pre-inoculum and grown at 37 °C and 210 rpm. At an OD_600nm_ of 0.8, IPTG was added to a final concentration of 0.5 mM and cells were grown at 25 °C and 120 rpm for 20 hours. The cells were harvested at 8739 *g*, 6 °C for 15 min and resuspended in 50 mM of Tris-HCl, pH 7.6.

### 2.2 Purification of AniA

Prior to cell lysis, a cocktail of protease inhibitors (cOmplete Mini, EDTA-free Protease inhibitors, Roche), DNase I (Roche) and 10 mM CuSO_4_ were added to the cell suspension. Cells were disrupted with a French Press by subjecting the cells to a pressure of 18000 psi. The lysed cells were centrifuged at 41399 *g* and 6 °C and for 1 hour (Beckman Avanti J-25 centrifuge) to obtain the cytoplasmic protein extract.

The purification protocol was adapted from the one described in [15] consisting of two purification steps. The cytoplasmic extract was applied onto an immobilized nickel-sepharose matrix (5 mL HisTrap HP, Cytiva) equilibrated with 20 mM Tris-HCl, pH 8.0. and 500 mM NaCl. AniA was eluted with a gradient of 0 to 500 mM of imidazole in the equilibration buffer. The fractions containing AniA were concentrated over an Amicon® Ultra-15 Centrifugal Filter Unit with 30 kDa MWCO (Millipore). The buffer was exchanged to 10 mM Tris-HCl, pH 7.6 using a Sephadex G-25 PD10 (Cytiva) desalting column, equilibrated in the same buffer. This AniA sample was then applied onto an anion exchange column (Resource Q, Cytiva) equilibrated with 20 mM Tris-HCl, pH 7.6. AniA was eluted with a gradient from 0 to 500 mM of NaCl in the equilibration buffer and the fractions containing the protein were concentrated and buffer exchanged to 10 mM Tris-HCl, pH 7.6 with an Amicon® Ultra-15 Centrifugal Filter Unit with 30 kDa MWCO (Millipore). The pure AniA was stored at −80 °C until further use. To analyze the purity of the samples collected during the purification process, 12.5 % SDS-PAGE in Tris-Tricine buffer was performed at 150 V for 70 minutes and the gel stained for the presence of protein with Coomassie Blue.

Protein and copper content were determined using the modified Lowry method [25], using bovine serum albumin (BSA) (Sigma) as the standard protein, and using a modified version of the copper(I)-biquinoline assay [26], respectively.

### 2.3 Spectroscopic and thermal characterization

UV-visible spectra were obtained in a Shimadzu UV-1800 spectrophotometer. The molar extinction coefficients were determined based on the copper concentration. The reduced and oxidized form of AniA were obtained by addition of sodium dithionite or potassium ferricyanide, respectively.

Electron Paramagnetic Resonance (EPR) was performed in a Bruker EMX spectrometer (Bruker BioSpin GmbH, Germany) coupled to an Oxford Instruments ESR-900 (Oxford Cryosystems, UK) continuous flow helium cryostat and a high-sensitivity perpendicular-mode rectangular cavity. The EPR spectra were acquired at a microwave frequency of 9.40 GHz, a microwave power of 0.202 mW, modulation amplitude of 0.5 mT and receiver gain of 1.0 × 10^5^. Other experimental conditions are listed in the figure legend.

Circular Dichroism (CD) in the far-UV and visible regions were performed in an Applied Photophysics ChirascanTM qCD spectrometer. AniA was prepared in 20 mM sodium phosphate, pH 7.0 at 2.7 μM and 105 μM for the far-UV and visible regions, respectively. In the far-UV region, spectra were obtained prior, during and after a temperature ramp that varied between 10 °C and 90 °C with an increase of 2 °C/min and 1 min stabilization. The far-UV data analysis for secondary structure determination was performed with the servers BeSTSel [27, 28] and Dichroweb [29, 30]. The characterization of AniA’s unfolding process was analyzed with the Van’t Hoff Plot, as described in [31], considering a two-state equilibrium between the folded native state and the unfolded state.

Differential Scanning Calorimetry (DSC) was performed using a TA™ Nano DSC calorimeter and the data was analyzed with NanoAnalyze™ software. AniA at 15 μM was prepared in 10 mM Hepes, pH 7.5, and the blank sample was the sample buffer.

### 2.4 Re-oxidation of AniA in the presence of nitrite

Re-oxidation of AniA in the presence of nitrite was assessed by UV-visible spectroscopy. As-isolated AniA (18.5 μM) was initially reduced with either 2 mM of sodium dithionite or 1 mM of sodium ascorbate, in the latter the reduction was mediated with 5 μM of diaminodurol and AniA was incubated for 30 minutes. Then, 5 mM sodium nitrite was added to the reduced AniA and spectra were acquired for up to 30 minutes of incubation with the substrate. The re-oxidation of AniA after sodium nitrite addition was assessed at pH 6.0 and pH 7.4 in 20 mM sodium phosphate and 9.5 in 20 mM of 3-(cyclohexylamino)-1-propanesulfonic acid (CAPS).

## 3. Results and Discussion

### 3.1 Protein production

The soluble domain of AniA was heterologously produced in *E. coli* as a cytoplasmic 36.4 kDa protein (trimer with 109 kDa), as a signal peptide is absent in the construct used (Figure S1, in Supplementary Materials). The enzyme was purified in two chromatographic steps: an affinity chromatography (as it has a His-tag at the C-terminus) followed by an anionic chromatography (as AniA’s pI is 5.7, considering its primary sequence). The protein was judged pure by its SDS-PAGE (Figure 2A) and had a purity ratio of A_598nm_ of the oxidized state to A_277nm_ of as-isolated state of ∼ 0.1. This purification procedure yielded 15-30 mg pure AniA/L of culture medium.

**Figure 2.**
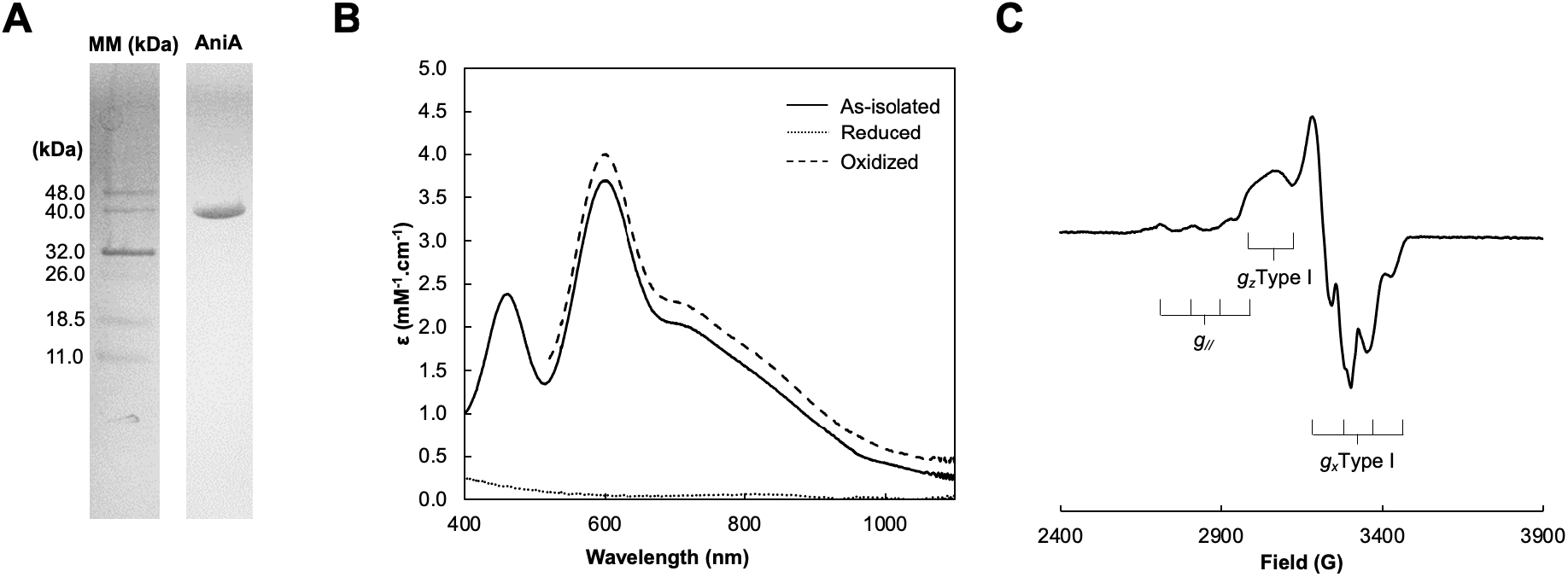
(**A**) Profile of pure AniA in a 12.5 % SDS-PAGE stained with Coomassie blue. Legend: MM – Molecular Marker, AniA – purified AniA. (**B**) Visible spectra of AniA in 10 mM Tris-HCl, pH 7.6. Legend: dashed, solid, and dotted line represent the ferricyanide-oxidized, as-isolated; as dithionite-reduced spectrum, respectively. (**C**) X-band EPR spectrum of 310 μM AniA in 10 mM Tris-HCl, pH 7.6 acquired at 30 K.

### 3.2 Spectroscopic characterization

#### UV-visible spectroscopy

The visible spectrum of AniA is presented in Figure 2B. The as-isolated AniA has the typical features of CuNiRs with two absorption bands at 458 nm and 598 nm with A_458nm_/A_598nm_ of 0.7, as previously reported [15]. In the oxidized spectrum, by addition of excess of sodium ferricyanide, the band at 598 nm slightly increases in intensity, whereas in the presence of excess sodium dithionite the absorption bands completely disappear. As these bands arise from transitions of T1Cu in the oxidized state, is it concluded that the as-isolated AniA has its T1Cu center only partially reduced.

The copper content per total protein was determined to be 1.8 ± 0.2, confirming that AniA was produced with both copper centers occupied. The copper quantification was used to determine the molar extinction coefficients of 2.76 mM^-1^.cm^-1^ and 4.0 mM^-1^.cm^-1^ at 458 nm and 598 nm, respectively.

T1Cu centers from blue copper proteins with two histidine, one cysteine and one methionine as coordinating residues typically have a distorted tetrahedral geometry [32]. As previously mentioned, the spectral features of CuNiRs arise from charge transfer transitions between the copper atom of the type-1 center and the sulfur of its coordinating cysteine. These transitions are directly influenced by the bond length between the copper atom and the sulfur of its coordinating cysteine and methionine residues that is dependent on the center’s geometry. With the increase of Cys(S)-Cu and decrease of Met(S)-Cu bonds, copper proteins are known to shift from blue to green color due to its absorption spectrum. Based on its resolved crystallographic structure, AniA has Met(S)-Cu of ∼ 2.5 Å and Cys(S)-Cu of ∼ 2.0 Å. Plastocyanin, a typical blue T1Cu protein, has a Met(S)-Cu ∼ 2.82 Å and Cys(S)-Cu ∼ 2.1 Å, whereas the green CuNir form *Achromobacter cyclocastes* (*Ac*NiR) has Met(S)-Cu ∼ 2.55 Å and Cys(S)-Cu ∼ 2.2 Å [18], confirming the direct influence of these bonds on the center’s geometry and subsequently spectroscopic features. Thus, considering these distances, AniA could be considered as part of the green CuNiRs class. However, in this class of proteins the ratio between both absorption bands is ≥ 1, as is the case of *Ac*NiR with A_464nm_/A_590nm_ of 1.3. Therefore, AniA cannot be fully considered as a green CuNiR.

The absorption bands of AniA have molar extinction coefficients closer to the ones reported for bluish-green CuNiR from *Bradyrhizobium japonicum*: 2.6 mM^-1^.cm^-1^ (458 nm) and 4.4 mM^-1^.cm^-1^ (592 nm) with a ε_458nm_/ε_592nm_ of 0.59 [34], then to the blue PsNiR from *Pseudomonas chlororaphis* with 9.87 mM^-1^.cm^-1^ (598 nm) [33] or to the green AcNiR with 3.9 mM^-1^.cm^-1^ (464 nm) and 3.0 mM^-1^.cm^-1^ (590 nm). Therefore, based on its UV-visible features AniA is considered to be a bluish-green CuNiR.

#### Electron Paramagnetic Resonance

The 30-K EPR spectrum of AniA has features of the presence of T1Cu and T2Cu centers (Figure 2C). T2Cu center has the typical features of an axial signal (*g*_//_ = 2.39 and A_//_ = 9.1 mT) similar to the one of *Achromobacter cycloclastes* CuNiR. T1Ccu center has a rhombic signal, with a small hyperfine coupling constant giving rise to an unresolved signal in the *g*_z_ region (*g*_*z*_ = 2.21). T1Cu signal has *g*_z_ = 2.27 (A_z_= 2.0 mT), *g*y= 2.14 (A_y_= 3.0 mT) and *g*_x_= 2.10 (A_x_ = 5.0 mT), with g-values similar to pseudoazurin [35], stellacyanin [36] and cucumber basic protein [37]. Belonging to the bluish-green class of CuNiRs, AniA EPR spectrum differs from normal blue and green copper sites. Although the absorbance at 598 nm is more intense than at 458 nm, the latter arises from a S(σ)cys→Cu strong enough that T1Cu shows a rhombic signal in EPR spectroscopy, contrary to what is seen in blue NiRs. A bluish-green CuNiR from *Bradyrhizobium japonicum*, with similar UV-vis spectrum [34], presents a different EPR spectrum, with nearly axial signals attributed to both T1Cu and T2Cu, although the g-values for g_z_ of both centers are similar to the ones observed in AniA. Thus, AniA combines the typical rhombicity of T1Cu signals from green copper centers, together with small hyperfine constants typical of blue copper centers.

#### Circular Dichroism

The secondary structure of the isolated AniA was examined by circular dichroism in the far-UV region. This spectrum has typical features of a predominant β-sheet structure, with a positive peak at 200 nm and a negative peak at 217 nm (Figure 3A). Green fluorescence protein [38] and intestinal fatty acid binding protein [39, 40] are known β-sheet proteins and have a similar far-UV spectra to AniA, thus confirming its β-sheet nature

**Figure 3.**
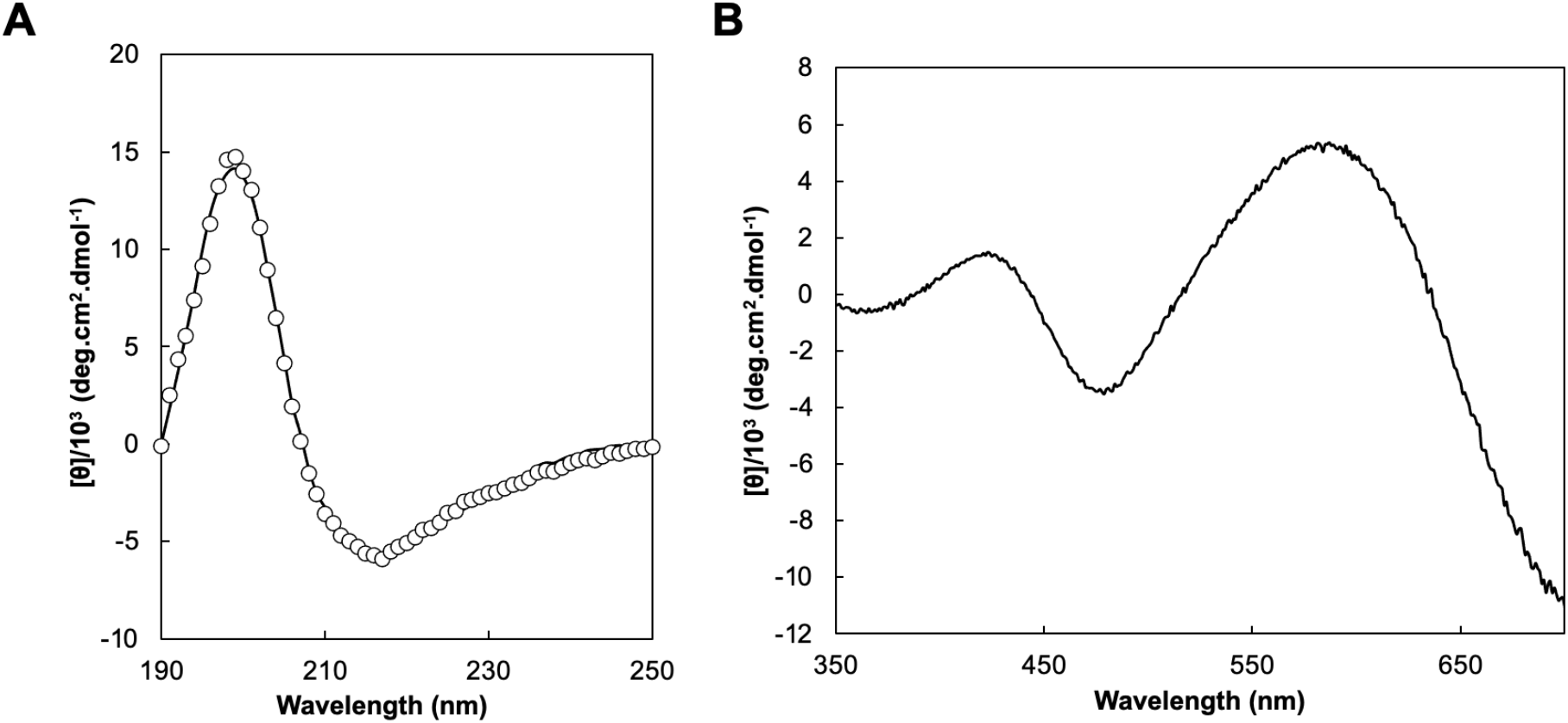
Circular dichroism spectra of AniA in the far-UV (**A**) and visible (**B**) regions. The spectra were obtained in 20 mM sodium phosphate pH 7.0 at 20 °C with 2.7 μM AniA for the far-UV and 105 μM AniA for the spectra in visible region.

The estimation of the secondary structure was performed with the servers BeStSel [27, 28] and DichroWeb [29, 30] and are presented in Table 1. The servers’ analysis of AniA’s far-UV spectrum determined 34-44 % of β-sheets, 3-12 % of α-helices, 11-25 % of turns and 17-47 % of unordered elements. There are some discrepancies between the different algorithms, depending on the secondary structure element, but overall agree with the values obtained from AniA X-ray structure to be mainly composed by β-sheets.

**Table 1.** Secondary structure elements of AniA determined from the X-ray structure and circular dichroism data. X-ray structure data retrieved from PDB:1KBW and data was simulated with BeStSel and DichroWeb - SELCON3, CONTIN, CSSTR and k2D programmes.

| Secondary Structure<br>Element | AniA<br>PDB ID 1KBW | BeStSel | Dichroweb |  |  |  |
| --- | --- | --- | --- | --- | --- | --- |
|  |  |  | SELCON3 | CONTIN | CSSTR | k2D |
| α-helix | 8 % | 3 % | 11 % | 11 % | 12 % | 12 % |
| β-sheet | 50 % | 44 % | 42 % | 48 % | 34 % | 41 % |
| Turn | 12 % | 11 % | 20 % | 25 % | 25 % | - |
| Unordered | 29 % | 42 % | 27 % | 17 % | 29 % | 47 % |

The CD spectrum in the visible region has two main positive peaks at 425 nm and 585 nm, that resemble the ones observed in the UV-visible spectrum, and a minimum that seems to form at around 700-800 nm, although data was not collected at higher wavelengths to confirm it. This peak with negative ellipticity is due to d-d weak transitions, also detectable in visible spectrum. As CD spectrum in the visible region arises from prosthetic groups, the coordination and geometry of the type-1 copper center and its neighbouring residues will influence the observed peaks [41]. AniA’s visible CD spectrum is similar to the one reported for *Hyphomicrobium* sp. A3151 CuNiR [42], but different from *Paracoccus pantotrophus* type-1 copper pseudoazurin, in which the S(π)cys→Cu charge transfer band appears as a peak with negative ellipticity [35]. Furthermore, CuNiR M150Q variant of *Ac*NiR from *Achromobacter cycloclastes* IAM1013 has differences in the visible CD spectrum compared to the wild-type [43]. The changes observed in M150Q-*Ac*NiR spectrum were attributed to a distortion in the geometry, leading to changes in the CD spectrum, and by the loss of chirality in one of the ligands. Therefore, the observed peaks in AniA visible CD are not only due to the S(σ)cys→Cu (425 nm) and S(π)cys→Cu (585 nm) charge transfer transitions, but also influenced by the distorted tetrahedral geometry and chirality of the ligands, especially the axial methionine.

### 3.3 Thermostability

Both CD and DSC were used to access the protein thermostability. Thermal denaturation of AniA was monitored by CD in the far-UV and visible regions (Figure 4, Panels A and C). CD spectra were acquired for each temperature, in the 10 ºC – 90 ºC range, whereas in DSC the energy of the unfolding process was followed alongside the temperature ramp up to 110 ºC (Figure 5).

**Figure 4.**
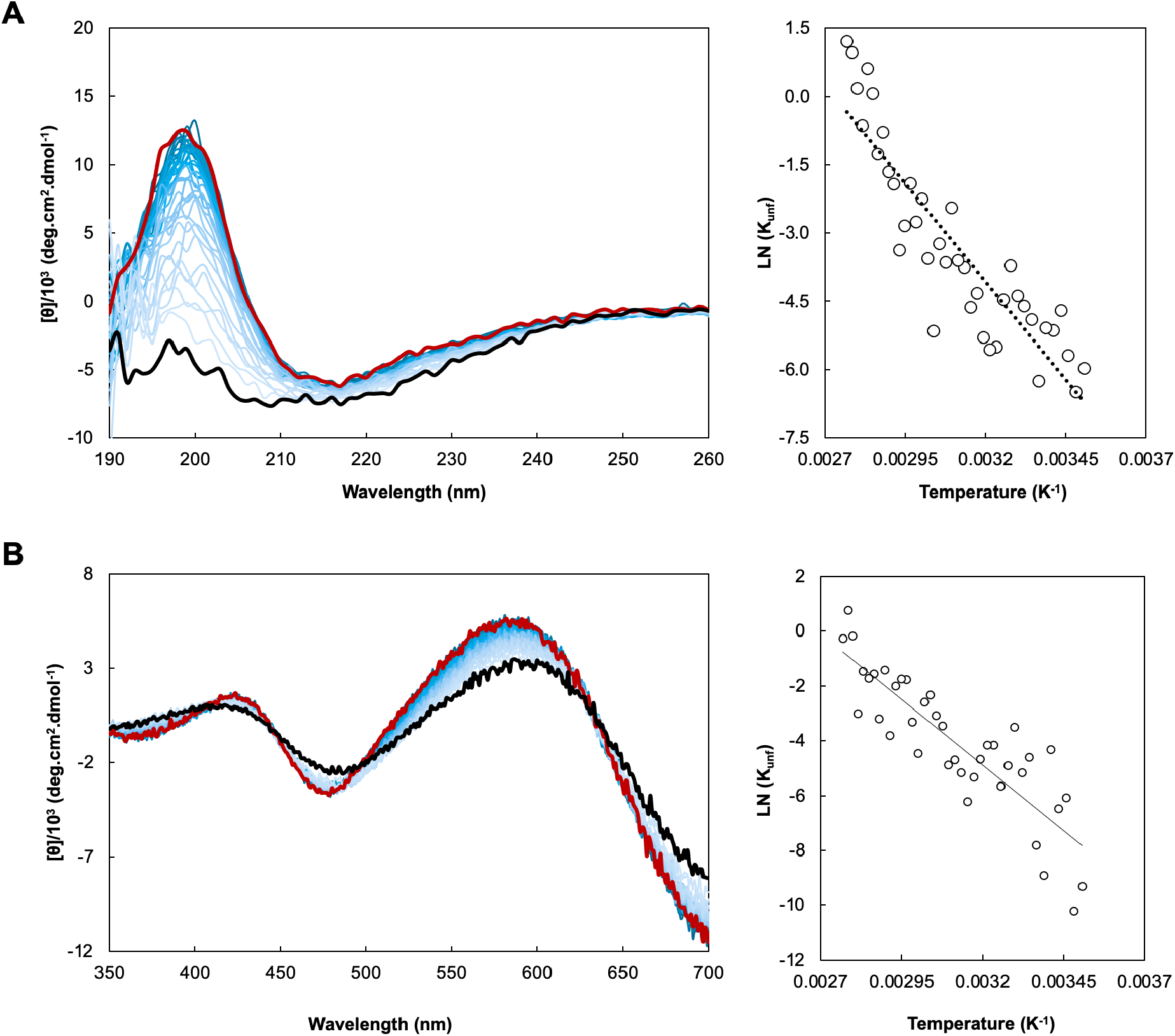
Thermal denaturation of AniA monitored by circular dichroism in the far-UV (top) and visible (bottom) regions. (**A**) and (**C**), spectra acquired during a temperature ramp between 10 ºC (red) and 90 ºC (black). (**B**) and (**D**), Van’t Hoff representation of ellipticity’s temperature variation at 196 nm (far-UV) and 598 nm (visible) (linear regression y=-8644.1x+23.585, R^2^=0.80 and y=-9582.9x+25.773, R^2^=0.74, respectively).

**Figure 5.**
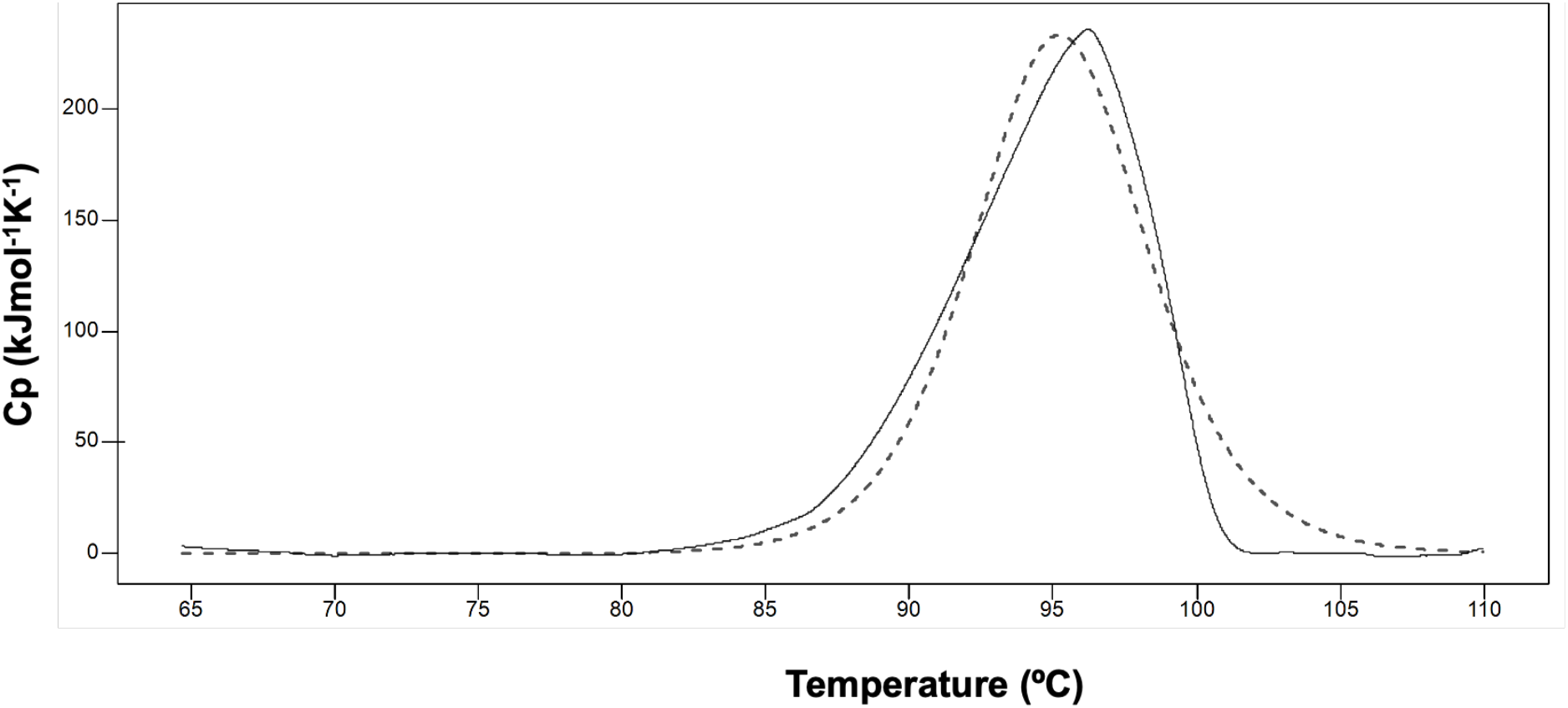
Thermogram of AniA in 10 mM Hepes, pH 7.5. Solid line is the experimental data, and dashed line the fitted thermogram with the parameters mentioned in the text.

AniA seems to be very stable, as the protein only starts to unfold and partially lose its structure at high temperatures. Assuming that AniA has two states – native and denatured, and only one transition between these states, the Van’t Hoff representations [44] (at 196 nm and 598 nm for UV and visible regions, respectively) allowed the estimation of thermostability parameters (Figure 4, Panels B and D). For the far-UV region, the enthalpy (ΔH_v_), entropy (ΔS_v_) and melting temperature (T_M_) were estimated to be ΔH_v_ = 72 ± 6 kJ/mol, ΔS_v_ = 0.20 ± 0.02 kJ/mol.K and T_M_ = 93 ± 3 ºC. For the visible region, the estimated parameters were ΔH_v_ = 80 ± 8 kJ/mol, ΔS_v_ = 0.20 ± 0.02 kJ/mol.K and T_M_ = 99 ± 3 ºC. Moreover, the CD spectra in the visible region did not disappear at 90 ºC, being observed a shift in the positive peaks. This indicates that the T1Cu coordination sphere is maintained to some degree up to this temperature, which agrees with the EPR spectra obtained at different temperatures for *Alcaligenes faecalis S-6* NiR [45], showing features of the presence of this center up to 90 ºC.

The thermogram obtained with DSC shows a single endothermic peak, from which a T_M_ of 95.3 ºC was estimated, as well as a ΔH_cal_ of 559.9 kJ/mol and an Aw of 3.3, indicating that the protein is in the trimeric form.

The unfolding temperatures obtained from CD and DSC indicate that AniA is in fact a very stable protein, which is corroborated by its ΔH_cal_. The T_M_ of AniA is similar to *Alcaligenes faecalis S-6* NiR [45], with a T_M_ of 99.6 ºC, which also has a high ΔH_cal_ value (1634 kJ/mol). This high thermostability might reflect the fact that AniA can exist extracellularly, and can require not losing the ability to reduce nitrite during infection [46].

### 3.4 Re-oxidation of AniA in the presence of nitrite

The re-oxidation of AniA in the presence of nitrite was assessed by visible spectroscopy, for the enzyme reduced with dithionite or ascorbate. These studies monitored the re-oxidation of T1Cu upon addition of nitrite in the presence of excess of reducing agent (5:1), by analyzing the increase in intensity of the absorption bands at 598 nm and 458 nm. These assays were performed at pH 6.0, 7.4 and 9.5 as described in Section 2.4 of Materials and Methods.

In Figure 6 Panel A represents the spectra of the assay with dithionite-reduced AniA, whereas in panel B are the spectra obtained with the ascorbate-reduced AniA, at the three tested pH values.

**Figure 6.**
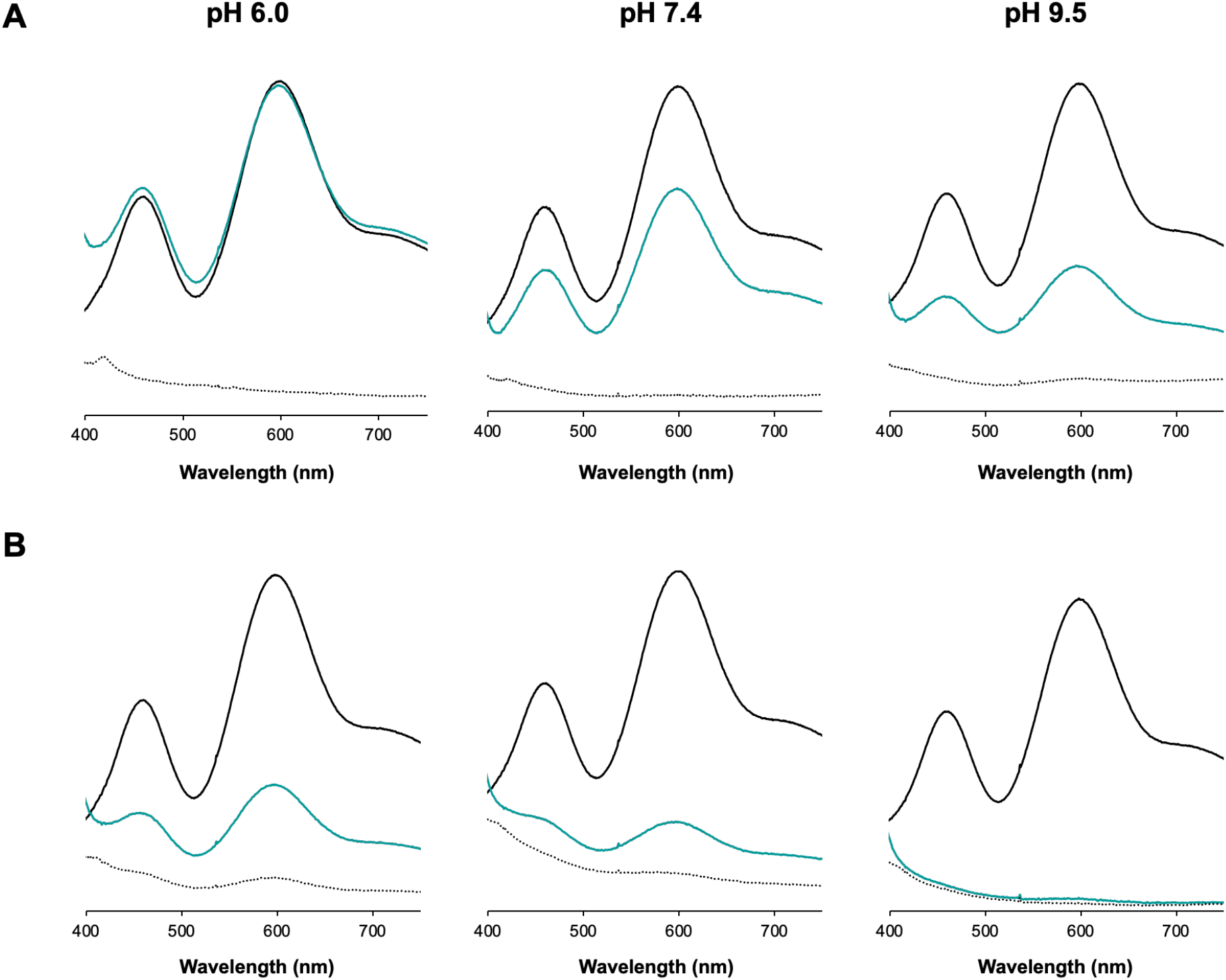
Reoxidation of dithionite (**A**) and ascorbate-reduced (**B**) AniA upon addition of sodium nitrite at pH 6.0 (left panel), 7.4 (middle panel) and 9.5 (right panel). Black solid spectra, as-isolated AniA; dotted black spectra – dithionite-reduced AniA; blue spectra - dithionite-reduced AniA upon nitrite addition after 5 minutes of incubation.

The spectra of the as-isolated AniA at the three tested pH values completely overlap (see Figure S2 in Supplementary Materials). As reported in the literature, charge transfer transitions (CT) of T1Cu are responsible for the visible spectrum of CuNiRs, with the absorption band at 598 nm arising from a CT of S(Cys) π → Cu 3dx2→y2 and the absorption band at 458 nm from a CT of S(Cys) σ → Cu 3dx2→y2. Therefore, any change of the copper center’s geometry would influence these transitions and subsequently be reflected in AniA’s visible features. As A_458nm_/A_598nm_ of the as-isolated spectrum remained unaltered at the different pH values, the geometry of T1Cu is not affected by pH.

The addition of both reducing agents led to the complete disappearance of AniA’s T1Cu absorption bands, confirming the reduction of this center. Upon addition of nitrite, both bands reappear, which is indicative of T1Cu re-oxidation due to electron transfer to T2Cu for nitrite reduction. The re-oxidation rate for both dithionite- and ascorbate-reduced AniA is higher at pH 6.0, with a re-oxidation of T1Cu after 5 minutes of incubation with nitrite of 99 % and 38 % of the initial intensity, respectively. For the other pHs there is marked decrease in the amount of re-oxidized AniA. Similar results were reported for *Bradyrhizobium japonicum* CuNiR [34]. In fact, a higher activity at acidic pH is expected as nitrite reduction required protons.

The re-oxidation rates are lower for the ascorbate-reduced AniA, which can be related with the reducing power of these reducing agents used.

## 4. Conclusions

This work presents the first spectroscopic characterization of AniA, the copper nitrite reductase from the pathogen *N. gonorrhoeae*. The recombinant AniA, solely comprising the soluble domain, was successfully isolated with both copper centers occupied. The UV-visible spectrum has the typical absorption bands at 598 nm and 458 nm with A_458nm_/A_598nm_ of 0.7, as previously reported [15]. These absorption bands are characteristic of green CuNiRs typically with A_460_/A_600_ above 1, conversely to what is observed in AniA. In addition to UV-visible spectroscopy, AniA EPR spectrum also has features of both green and blue CuNiRs, such as the rhombic nature of T1Cu EPR signal of green CuNiRs and the small hyperfine constants of blue CuNiRs. Therefore, is it confirmed that AniA is a bluish-green CuNiR. The hyperthermophilic nature of AniA, studied by DSC and CD (far-UV and visible regions), and higher reoxidation rates in the presence of nitrite at acidic pH, analyzed by visible spectroscopy, might be related with its extracellular location and the environment of its infection site.

## Author contribution

DB and RO performed the experiments with contributions from SRP, analyzed the data and wrote the manuscript. SRP planned the manuscript and experiments, performed, and analyzed data and wrote the manuscript.

## Acknowledgments

This work was financed by national funds from FCT - Fundação para a Ciência e a Tecnologia, I.P., in the scope of the project UIDP/04378/2020 and UIDB/04378/2020 of the Research Unit on Applied Molecular Biosciences - UCIBIO and the project LA/P/0140/2020 of the Associate Laboratory Institute for Health and Bioeconomy - i4HB. FCT supported SRP through the projects PTDC/BIA-PRO/109796/2009 and PTDC/BIA-BQM/29442/2017, DSB and RNSO through the scholarships UI/BD/151168/2021, and “Verão com Ciência2020”, respectively. We also thanks Dr. Filipe Folgosa and Prof. Miguel Teixeira for the assistance in the acquisition of EPR data.

## References

[1] W.G. Zumft, Cell biology and molecular basis of denitrification, Microbiol Mol Biol Rev 61 (1997) 533–616

[2] J.W. Allen, P.D. Barker, O. Daltrop, J.M. Stevens, E.J. Tomlinson, N. Sinha, Y. Sambongi, S.J. Ferguson, Why isn’t ‘standard’ heme good enough for c-type and d1-type cytochromes?, Dalton Trans (2005) 3410–3418

[3] F. Cutruzzolà, M. Arese, G. Ranghino, G. van Pouderoyen, G. Canters, M. Brunori, Pseudomonas aeruginosa cytochrome C551: probing the role of the hydrophobic patch in electron transfer, J Inorg Biochem 88 (2002) 353–361

[4] I.V. Pearson, M.D. Page, R.J. van Spanning, S.J. Ferguson, A mutant of Paracoccus denitrificans with disrupted genes coding for cytochrome c550 and pseudoazurin establishes these two proteins as the in vivo electron donors to cytochrome cd1 nitrite reductase, J Bacteriol 185 (2003) 6308–6315

[5] Y. Fukuda, K.M. Tse, T. Nakane, T. Nakatsu, M. Suzuki, M. Sugahara, S. Inoue, T. Masuda, F. Yumoto, N. Matsugaki, E. Nango, K. Tono, Y. Joti, T. Kameshima, C. Song, T. Hatsui, M. Yabashi, O. Nureki, M.E. Murphy, T. Inoue, S. Iwata, E. Mizohata, Redox-coupled proton transfer mechanism in nitrite reductase revealed by femtosecond crystallography, Proc Natl Acad Sci U S A 113 (2016) 2928–2933

[6] N.G. Leferink, C. Han, S.V. Antonyuk, D.J. Heyes, S.E. Rigby, M.A. Hough, R.R. Eady, N.S. Scrutton, S.S. Hasnain, Proton-coupled electron transfer in the catalytic cycle of Alcaligenes xylosoxidans copper-dependent nitrite reductase, Biochemistry 50 (2011) 4121–4131

[7] E. Libby, B.A. Averill, Evidence that the Type 2 copper centers are the site of nitrite reduction by Achromobacter cycloclastes nitrite reductase, Biochem Biophys Res Comm 187 (1992) 1529–1535

[8] L.M. Murphy, F.E. Dodd, F.K. Yousafzai, R.R. Eady, S.S. Hasnain, Electron donation between copper containing nitrite reductases and cupredoxins: the nature of protein-protein interaction in complex formation, J Mol Biol 315 (2002) 859–871

[9] S. Suzuki, T. Kohzuma, Deligeer, K. Yamaguchi, N. Nakamura, S. Shidara, K. Kobayashi, S. Tagawa, Pulse Radiolysis Studies on Nitrite Reductase from Achromobacter cycloclastes IAM 1013: Evidence for Intramolecular Electron Transfer from Type 1 Cu to Type 2 Cu, J Am Chem Soc 116 (2002) 11145–11146

[10] R.R. Eady, S. Samar Hasnain, New horizons in structure-function studies of copper nitrite reductase, Coordination Chemistry Reviews 460 (2022)

[11] E.I. Solomon, R.K. Szilagyi, S. DeBeer George, L. Basumallick, Electronic structures of metal sites in proteins and models: contributions to function in blue copper proteins, Chem Rev 104 (2004) 419–458

[12] S. Horrell, D. Kekilli, R.W. Strange, M.A. Hough, Recent structural insights into the function of copper nitrite reductases, Metallomics 9 (2017) 1470–1482

[13] M.J. Boulanger, M. Kukimoto, M. Nishiyama, S. Horinouchi, M.E. Murphy, Catalytic roles for two water bridged residues (Asp-98 and His-255) in the active site of copper-containing nitrite reductase, J Biol Chem 275 (2000) 23957–23964

[14] K. Kataoka, H. Furusawa, K. Takagi, K. Yamaguchi, S. Suzuki, Functional analysis of conserved aspartate and histidine residues located around the type 2 copper site of copper-containing nitrite reductase, J Biochem 127 (2000) 345–350

[15] M.J. Boulanger, M.E. Murphy, Crystal structure of the soluble domain of the major anaerobically induced outer membrane protein (AniA) from pathogenic Neisseria: a new class of copper-containing nitrite reductases, J Mol Biol 315 (2002) 1111–1127

[16] F.E. Dodd, J. Van Beeumen, R.R. Eady, S.S. Hasnain, X-ray structure of a blue-copper nitrite reductase in two crystal forms. The nature of the copper sites, mode of substrate binding and recognition by redox partner, J Mol Biol 282 (1998) 369–382

[17] R.W. Strange, L.M. Murphy, F.E. Dodd, Z.H. Abraham, R.R. Eady, B.E. Smith, S.S. Hasnain, Structural and kinetic evidence for an ordered mechanism of copper nitrite reductase, J Mol Biol 287 (1999) 1001–1009

[18] E.I. Solomon, Spectroscopic methods in bioinorganic chemistry: blue to green to red copper sites, Inorg Chem 45 (2006) 8012–8025

[19] N. Castiglione, S. Rinaldo, G. Giardina, V. Stelitano, F. Cutruzzola, Nitrite and nitrite reductases: from molecular mechanisms to significance in human health and disease, Antioxid Redox Signal 17 (2012) 684–716

[20] J.N. Sharma, A. Al-Omran, S.S. Parvathy, Role of nitric oxide in inflammatory diseases, Inflammopharmacology 15 (2007) 252–259

[21] P. Muenzner, C.R. Hauck, Neisseria gonorrhoeae Blocks Epithelial Exfoliation by Nitric-Oxide-Mediated Metabolic Cross Talk to Promote Colonization in Mice, Cell Host Microbe 27 (2020) 793–808 e795

[22] A.K. Mix, G. Goob, E. Sontowski, C.R. Hauck, Microscale communication between bacterial pathogens and the host epithelium, Genes Immun 22 (2021) 247–254

[23] X. Li, S. Parker, M. Deeudom, J.W. Moir, Tied down: tethering redox proteins to the outer membrane in Neisseria and other genera, Biochem Soc Trans 39 (2011) 1895–1899

[24] B.I. Baarda, R.A. Zielke, A.E. Jerse, A.E. Sikora, Lipid-Modified Azurin of Neisseria gonorrhoeae Is Not Surface Exposed and Does Not Interact With the Nitrite Reductase AniA, Front Microbiol 9 (2018) 2915

[25] G.L. Peterson, Meth Enzymol, vol. 91, Academic Press, 1983, pp. 95–119.

[26] S.R. Pauleta, F. Guerlesquin, C.F. Goodhew, B. Devreese, J. Van Beeumen, A.S. Pereira, I. Moura, G.W. Pettigrew, Paracoccus pantotrophus pseudoazurin is an electron donor to cytochrome c peroxidase, Biochemistry 43 (2004) 11214–11225

[27] A. Micsonai, F. Wien, L. Kernya, Y.H. Lee, Y. Goto, M. Refregiers, J. Kardos, Accurate secondary structure prediction and fold recognition for circular dichroism spectroscopy, Proc Natl Acad Sci U S A 112 (2015) E3095–3103

[28] A. Micsonai, F. Wien, E. Bulyaki, J. Kun, E. Moussong, Y.H. Lee, Y. Goto, M. Refregiers, J. Kardos, BeStSel: a web server for accurate protein secondary structure prediction and fold recognition from the circular dichroism spectra, Nucleic Acids Res 46 (2018) W315–W322

[29] A.J. Miles, S.G. Ramalli, B.A. Wallace, DichroWeb, a website for calculating protein secondary structure from circular dichroism spectroscopic data, Protein Sci 31 (2022) 37–46

[30] N. Sreerama, R.W. Woody, Estimation of protein secondary structure from circular dichroism spectra: comparison of CONTIN, SELCON, and CDSSTR methods with an expanded reference set, Anal Biochem 287 (2000) 252–260

[31] M.F. Anjum, T.M. Stevanin, R.C. Read, J.W. Moir, Nitric oxide metabolism in Neisseria meningitidis, J Bacteriol 184 (2002) 2987–2993

[32] J.T. Rubino, K.J. Franz, Coordination chemistry of copper proteins: how nature handles a toxic cargo for essential function, J Inorg Biochem 107 (2012) 129–143

[33] D. Pinho, S. Besson, C.D. Brondino, B. de Castro, I. Moura, Copper-containing nitrite reductase from Pseudomonas chlororaphis DSM 50135, Eur J Biochem 271 (2004) 2361–2369

[34] J.C. Cristaldi, F.M. Ferroni, A.B. Dure, C.S. Ramirez, S.D. Dalosto, A.C. Rizzi, P.J. Gonzalez, M.G. Rivas, C.D. Brondino, Heterologous production and functional characterization of Bradyrhizobium japonicum copper-containing nitrite reductase and its physiological redox partner cytochrome c550, Metallomics 12 (2020) 2084–2097

[35] X. Xie, R.G. Hadt, S.R. Pauleta, P.J. Gonzalez, S. Un, I. Moura, E.I. Solomon, A variable temperature spectroscopic study on Paracoccuspantotrophus pseudoazurin: protein constraints on the blue Cu site, J Inorg Biochem 103 (2009) 1307–1313

[36] A.A. Gewirth, S.L. Cohen, H.J. Schugar, E.I. Solomon, Spectroscopic and theoretical studies of the unusual EPR parameters of distorted tetrahedral cupric sites: correlations to x-ray spectral features of core levels, Inorg Chem 26 (1987) 1133–1146

[37] L.B. LaCroix, D.W. Randall, A.M. Nersissian, C.W.G. Hoitink, G.W. Canters, J.S. Valentine, E.I. Solomon, Spectroscopic and Geometric Variations in Perturbed Blue Copper Centers: Electronic Structures of Stellacyanin and Cucumber Basic Protein, J Am Chem Soc 120 (1998) 9621–9631

[38] N.V. Visser, M.A. Hink, J.W. Borst, G.N.M. van der Krogt, A.J.W.G. Visser, Circular dichroism spectroscopy of fluorescent proteins, FEBS Letters 521 (2002) 31–35

[39] N. Sreerama, S.Y. Venyaminov, R.W. Woody, Estimation of the number of alpha-helical and beta-strand segments in proteins using circular dichroism spectroscopy, Protein Sci 8 (1999) 370–380

[40] N. Sreerama, R.W. Woody, Structural composition of betaI- and betaII-proteins, Protein Sci 12 (2003) 384–388

[41] N.J. Greenfield, Meth Enzymol, vol. 383, Academic Press, 2004, pp. 282–317.

[42] S. Suzuki, T. Kohzuma, S. Shidara, K. Ohki, T. Aida, Novel spectroscopic aspects of type I copper in hyphomicrobium nitrite reductase, Inorganica Chim Acta 208 (1993) 107–109

[43] K. Kataoka, K. Yamaguchi, S. Sakai, K. Takagi, S. Suzuki, Characterization and function of Met150Gln mutant of copper-containing nitrite reductase from Achromobacter cycloclastes IAM1013, Biochem Biophys Res Commun 303 (2003) 519–524

[44] N.J. Greenfield, Using circular dichroism collected as a function of temperature to determine the thermodynamics of protein unfolding and binding interactions, Nat Protoc 1 (2006) 2527–2535

[45] A. Stirpe, R. Guzzi, H. Wijma, M.P. Verbeet, G.W. Canters, L. Sportelli, Calorimetric and spectroscopic investigations of the thermal denaturation of wild type nitrite reductase, Biochim Biophys Acta 1752 (2005) 47–55

[46] L.K. Shewell, S.C. Ku, B.L. Schulz, F.E. Jen, T.D. Mubaiwa, M.R. Ketterer, M.A. Apicella, M.P. Jennings, Recombinant truncated AniA of pathogenic Neisseria elicits a non-native immune response and functional blocking antibodies, Biochem Biophys Res Commun 431 (2013) 215–220

