## Supplementary figures and images for "Biochemical characterization of AniA from *Neisseria gonorrhoeae*"

### Figure S1

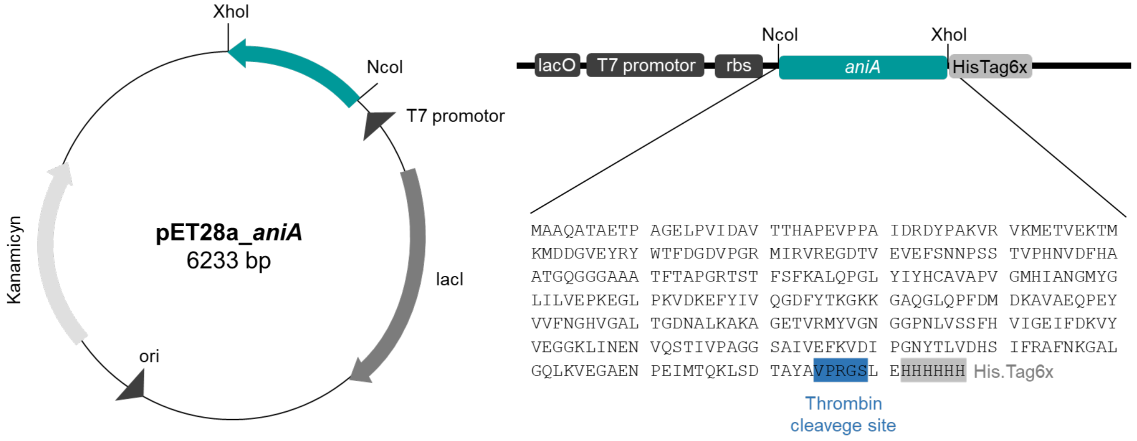


**Figure S1.** Schematic representation of AniA construct

### Figure S2

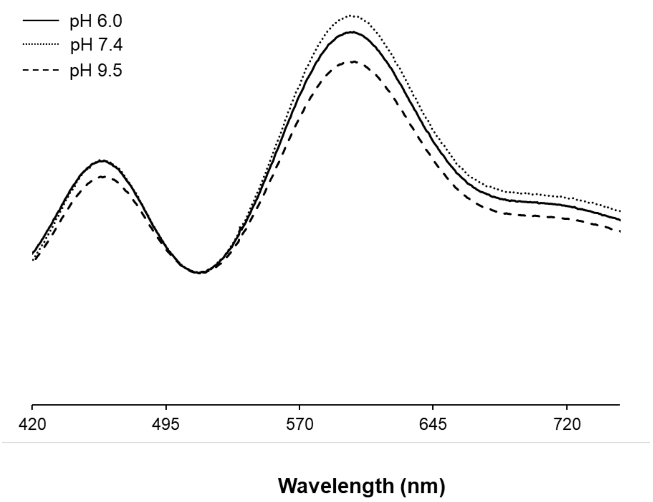


**Figure S2.** UV-visible spectra of as-isolated AniA (18.5 μ M) at pH 6.0 (full line), 7.4 (dotted line) and 9.5 (traced line).
